# A Bioengineered Platform for Enhanced Observability of Patterned Gastrointestinal Organoid Monolayers with Bilateral Access

**DOI:** 10.1101/2024.01.26.577381

**Authors:** Moritz Hofer, Maria A. Duque-Correa, Matthias P. Lutolf

**Author notes:** Corresponding author: Matthias P. Lutolf. Contact person regarding caecaloid model and whipworm infection: Maria A. Duque-Correa.

## Abstract

Studying the physiology and pathology of gastrointestinal (GI) tissues requires tools that can accurately mimic their complex architecture and functionality *in vitro*. Organoids have emerged as one such promising tool, though their closed structures with poorly accessible lumen and limited observability makes readouts challenging. In this study, we introduce a bioengineered organoid platform that generates bilaterally accessible 3D tissue models, allowing independent manipulation of both the apical and basal sides of patterned epithelial monolayers. We successfully constructed gastric, small intestinal, caecal, and colonic epithelial models that faithfully reproduce tissue-respective geometries and exhibit high physiological relevance, evidenced by the regionalization of stem cells and transcriptional resemblance to real epithelia. The gained observability allowed single-cell tracking over time and studies into the motility of cells in immersion and air-liquid interface cultures. Additionally, this model recapitulated *Trichuris muris* infection of the caecum epithelium, allowing the first live imaging of syncytial tunnel formation. Overall, this platform offers accessible organoids with improved observability, making it a valuable tool for investigating the dynamics of GI epithelial cells and their interactions with pathogens.

## Introduction

Studies into the physiology and pathology of the digestive system require *in vitro* models that accurately recapitulate the complex architecture and functionality of gastrointestinal (GI) tissues. Over the last decade, significant progress has been made in developing organoids, which are 3D culture systems derived from primary cells or pluripotent stem cells that provide a source of untransformed cells, including GI epithelial cells^1–3^. Such 3D organoids take advantage of the capacity of stem cells to self-renew and proliferate into diverse differentiated cell progeny when instructed with the necessary niche factors mimicking the native tissue environment. These cues include soluble growth factors and morphogens as well as attachment cues from the extracellular matrix (ECM) and its physical parameters. Epithelial organoids commonly grow to luminal structures that display self-organized geometries that partially resemble the *in vivo* architecture, exemplified by intestinal organoids which develop structures with remarkable resemblance to the intestinal crypt^4^. Cells in such organoids show apical-basal polarization with the apical side facing the lumen, which is very challenging to access. In consequence, organoids are as such little useful for studying important gastrointestinal processes that require experimental control of the apical side, such as exposure to nutrients and interactions with pathogens. Furthermore, organoids grow constantly and naturally shed dead cells accumulate in the lumen, which ultimately limits the organoid lifespan to only several days, preventing studying longer-term pathophysiological processes. In addition, due to the stochastic nature of the differentiation events involved in the growth and maturation of these organoids, they exhibit significant heterogeneity regarding their cellular composition as well as their size and shape, which ultimately makes optical as well as biochemical readouts very challenging^5^. In summary, the applicability of organoids is hampered by their poor observability, restricted accessibility for manipulation, and challenges in simulating long-term phenomena.

To circumvent these issues while retaining the advantages of having multiple cell types, organoids have been used as source of cells for 2D cultures. Grown on ECM-coated substrates, these 2D cultures can be kept for extended periods, and when using transwells, both sides of the epithelium can be easily manipulated^6–14^. However, even though clusters of stem cells and differentiated cells can arise in distinct regions of these 2D cultures, the corresponding geometry is missing and the patterning is seemingly stochastic. To address this lack of geometrical patterning, hydrogels with patterned surfaces have been employed as scaffolds. These approaches succeeded in shaping such organoid-derived monolayers to mimic the structural and cellular arrangement of native tissues^6,15–19^, but they either pose challenges for live microscopy or lack convenient experimental control.

In this study, we present a novel platform that combines the advantages of classical transwell systems with the benefits of organoids grown on hydrogel scaffolds. Our membrane-free platform enables the generation of bilaterally accessible 3D tissue models, facilitating independent manipulation of both the apical and basal sides of patterned epithelial monolayers and allowing excellent observability using microscopy techniques. Using organoid-derived cells, we constructed gastric, small intestinal, caecal, and colonic epithelial models reflecting the geometries of the tissues. We assessed the dynamics of gastric epithelia in air-liquid interface cultures and recapitulated key features of *Trichuris muris* infection in caecal epithelia.

## Results

### Generation of a transwell surrogate chip for stem cell-derived epithelial monolayer with bilateral accessibility

We aimed to develop a transwell surrogate platform that combines bilateral accessibility with enhanced observability. To achieve this, we designed a polydimethylsiloxane (PDMS) microfluidic chip with phase-guiding pillars to cast a thin layer of collagen-based hydrogel (**Fig. 1a,b, Supplementary Fig. S1a**), with the hydrogel serving as a cell-adhesive scaffold. As in previous studies, we supplemented the hydrogel with 25% basement membrane extract (Matrigel^™^) to support epithelial stem cell maintenance^18^. Epithelial cells, such as the gastric epithelial cells obtained from 3D organoids here, were seeded on the upper side of the hydrogel disc, where they formed a monolayer (**Fig. 1c**). The hydrogel also ensures separation between the two compartments containing medium, which can be controlled independently: the medium on the upper side of the cell monolayer (apical medium) is added to the upper well-shaped opening of the microfluidic chip, while the medium the cells encounter on their basal side (facing the hydrogel) diffuses from the reservoir through the hydrogel. The efficiency of medium diffusion was modeled and experimentally validated using 40 kDa FITC-dextran (**Fig. 1d,e**). Additionally, confocal z-stack imaging confirmed the segregation of two separate media, each supplemented in their respective compartments with a differently fluorescently labeled dextran (**Fig. 1f**).

**Fig. 1.**
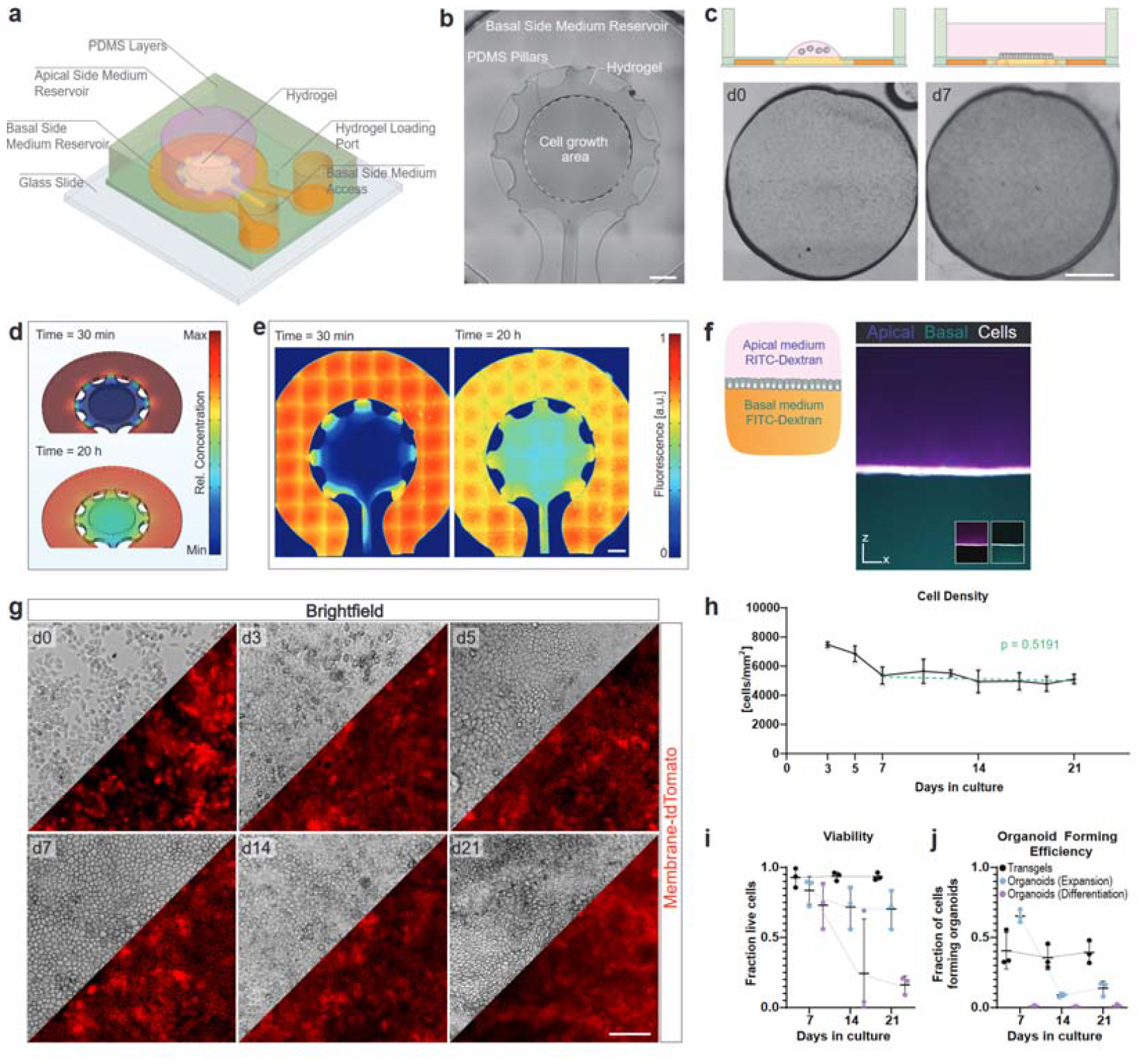
*Transgels:* A Microfluidic platform for stem-cell derived epithelial monolayer with bilateral accessibility. **a**, Schematic 3D representation of the microfluidic device made of two layers of PDMS for loading and casting a hydrogel scaffold for cell attachment and growth, with two distinct compartments for media for the apical and basal side. **b**, Microscopy overview image showing the hydrogel casted by PDMS pillars. Indication of accessible hydrogel surface intended for cell growth. Scale bar, 1 mm. **c**, Schematics and microscopy image of gastric epithelial cells seeded on the hydrogel opening and formation of monolayer. Scale bar, 1mm. **d,e**, Computational modeling (d) and experimental validation (e) of model growth factor diffusion from the basal medium compartment through the hydrogel. Scale bar, 1mm. Representative images of three independent experiments. **f**, Experimental confirmation of maintenance of two distinct medium compartment upon epithelial cell monolayer growth on hydrogel. Apical and basal side media were supplemented with RITC-, and FITC-labelled 40 kDa Dextran, respectively. Side projection of a confocal z-stack image of media and cells (mTmG, membrane-localized tdTomato), acquired 24 h after medium change. Inserts displaying overlay of cells with only one media, showing the complete absence of media in the other compartment, respectively. Representative image of three independent experiments. Scale bar, 100 μm. **g,h**, Brightfield and fluorescent images of mTmG gastric epithelial monolayer cultured with stem cell growth factors and morphogens (Wnt-3a, R-spondin, Noggin; “GF”) uniquely present in the basal side medium from day 3 on and (h) the quantification of cell density over time. Points represent mean and SD of three independent experiments. p-value represents significance of the fitted linear model (day 7-21) deviating from zero. Scale bar, 100 μm. **i,j**, Gastric epithelial cells were collected from Transgel hydrogels and analyzed for viability (i) and capacity to form organoids (j). Comparison with cells from 3D organoid culture in expansion (with GF) or differentiation (without GF). Mean and SD from three independent experiments are shown.

To demonstrate the functionality of the platform, we seeded gastric epithelial cells as before, but kept the growth factors and morphogens necessary for stem cell maintenance^20^ (Wnt3a, R-Spondin, and Noggin, collectively termed “GFs”) on the basal side only and replaced the apical medium with medium lacking GFs, once the monolayer was formed (day 3). Still, the monolayer could be maintained with consistent cell density for up to three weeks, indicating that delivery of GFs uniquely from the basal side is sufficient (**Fig. 1g,h, Supplementary Fig. S1b,c**). Importantly, during medium changes, apically shed dying cells were removed, resulting in high overall viability over time without the need of passaging, which contrasts with 3D organoids in expansion (with GFs) or differentiation (without GFs) medium, where dead cells accumulate in the lumen (**Fig. 1i, Supplementary Fig. S1d**). We collected the cells from the hydrogel surfaces at different time points and assessed their capacity to regenerate 3D organoids. A stable proportion of organoids were formed from cells derived from these monolayers, which could not be achieved with cells from traditional organoid cultures without passaging, indicating the continuous presence of stem cells in the monolayers (**Fig. 1j**). Considering the advantages offered by this novel platform, which combines features of classical transwell systems and hydrogel scaffolds, we term it *Transgel*.

### Observable gastrointestinal model systems using scaffold patterning

Hydrogel scaffolds have been generated previously for intestinal epithelial organoid cultures to mimic the crypt-villus architectures, which instruct the epithelial stem cells to generate cell type patterns reminiscent of the native tissues^6,15–19^. Similarly, we employed PDMS stamps to shape the surface of the hydrogel scaffolds in the Transgel devices to resemble the *in vivo* architecture of epithelial monolayers from various GI tissues (**Fig. 2a,b**)^21–23^. For stomach, caecum, and colon tissues, we employed an array of crypts or glands of different depths, while for the small intestine, we included a villus domain. Cells derived from 3D organoids of the respective tissues were seeded and allowed to colonize the surface for three days. *In vivo*, the stem-cell populations in the glands and crypts are sustained by GFs secreted by the underlying mesenchyme^21^. To recapitulate this situation in our *in vitro* models, we reduced the concentration of GFs on the apical side after three days in culture, allowing the cells to differentiate for an additional two to four days, depending on the tissue model (**Supplementary Fig. S2a**). After this differentiation, the cellular monolayer retained the predetermined structure dictated by the scaffold. We used live confocal z-stack imaging of cell membranes using mTmG cells to highlight the excellent observability of the generated model systems, enabling the detection of single cells and the observation of their individual shapes (**Fig. 2c, Supplementary Video 1**). All model epithelia exhibited apical-basal polarity, with the unilateral accumulation of F-actin and Itga6, respectively (**Supplementary Fig. S2b**).

**Fig. 2.**
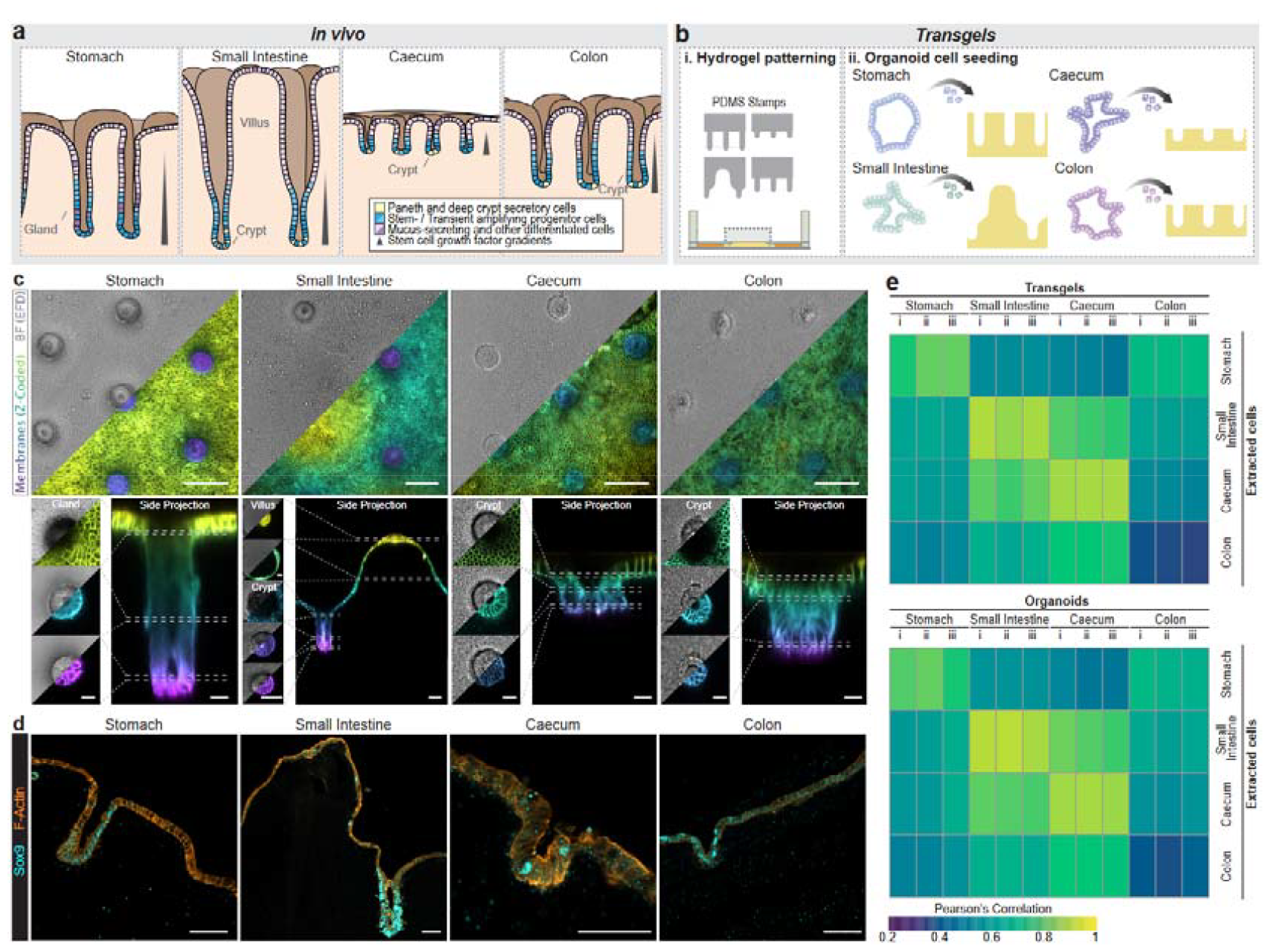
Generation of observable epithelial organoid monolayer of gastrointestinal tissues. **a**, Schematics of gastrointestinal epithelia. Stomach, caecum and colon regions display a gland/crypt-architecture and the small intestinal epithelia consists additionally of villi. Stem cells reside at the bottom of the invaginations. **b**, Setup of Transgel organoids. Hydrogel surfaces are geometrically patterned with polydimethylsiloxane (PDMS) stamps where subsequently cells of organoids are seeded onto. **c**, Fully grown Transgel organoids after 3 days of expansion and additional 4 days (stomach) or 2 days (others) of differentiation with reduced growth factors on apical side (see methods). mTmG cells expressing membrane-localized tdTomato were used. Extended field of depth (EDF) and maximum z-projection of z-colored membranes images of live confocal microscopy. Bottom: Single z slices and side projections. Scale bars, 100 μm (top panels), 50 μm (bottom panel for small intestine), 20 μm (bottom panels for other tissues). **d**, Immunohistochemical staining for Sox9 on sections of organoids grown in Transgels. Scale bars, 50 μm. **e**, Correlation score of the transcriptome (via RNA sequencing) of Transgel organoid cultures (top) and organoids grown in 3D (bottom) compared to that of freshly extracted epithelial cells. The mean expression values of thee independent extraction were used for freshly extracted cells.

To assess whether the constructed tissue models recapitulate the characteristic restriction of stem cells to the crypt and gland regions, we performed immunostaining on cryosections to mark SRY-box transcription factor 9 (Sox9)-positive stem- and progenitor cells. We found that Sox9-positive cells predominantly, but not exclusively, located in the crypt/gland regions across all tissue models (**Fig. 2d**). Similarly, staining for Ki-67 revealed higher proliferative activity in cells within the crypt/gland regions compared to the surface regions (**Supplementary Fig. S2c**). We also observed apical layers positive for glycoproteins stained with Alcian blue, suggesting the presence of mucus-secreting cells in all tissue models (**Supplementary Fig. S2d**). To comprehensively compare our new tissue models with traditional 3D organoids and freshly extracted epithelial cells (**Supplementary Fig. S3a**), we analyzed the transcriptome using RNA sequencing. Focusing on signature genes of the tissues, the Transgel organoid models correlated well with their *in vivo* tissue counterparts and were very similar to 3D organoids, indicating the preserved physiological relevance of 3D organoids in our new engineered organoids (**Fig. 2e, Supplementary Fig. S3b,c**). In conclusion, we generated membrane-free bilaterally accessible gastrointestinal epithelial models with tissue-relevant geometries enforcing the recapitulation of the tissue-characteristic stem cell regions.

### Air-liquid interface culture of gastric tissue model enhances transcriptional similarity to real tissue

To promote cell differentiation and improve resemblance to real tissues, thereby enhancing physiological relevance, conventional transwell-based stomach epithelial models have employed air-liquid interface (ALI) cultures^7,24^. To understand the cellular behavior in response to air exposure and the absence of medium on the apical side, we established an ALI culture on gastric Transgel organoids 3 days after seeding (**Fig. 3a**). After four days in ALI culture, we observed the cells in the glands were thinner than cells in immersion (IMM) conditions (**Fig. 3b-d**). Immunofluorescent staining of sections revealed that in ALI cultures the stem-cell compartments were preserved, similar to the standard model in IMM conditions (**Supplementary Fig. S4a**). As expected, secreted mucus accumulated on the apical side, providing a physical barrier between the cells and the air (**Supplementary Fig. S4b**).

**Fig. 3.**
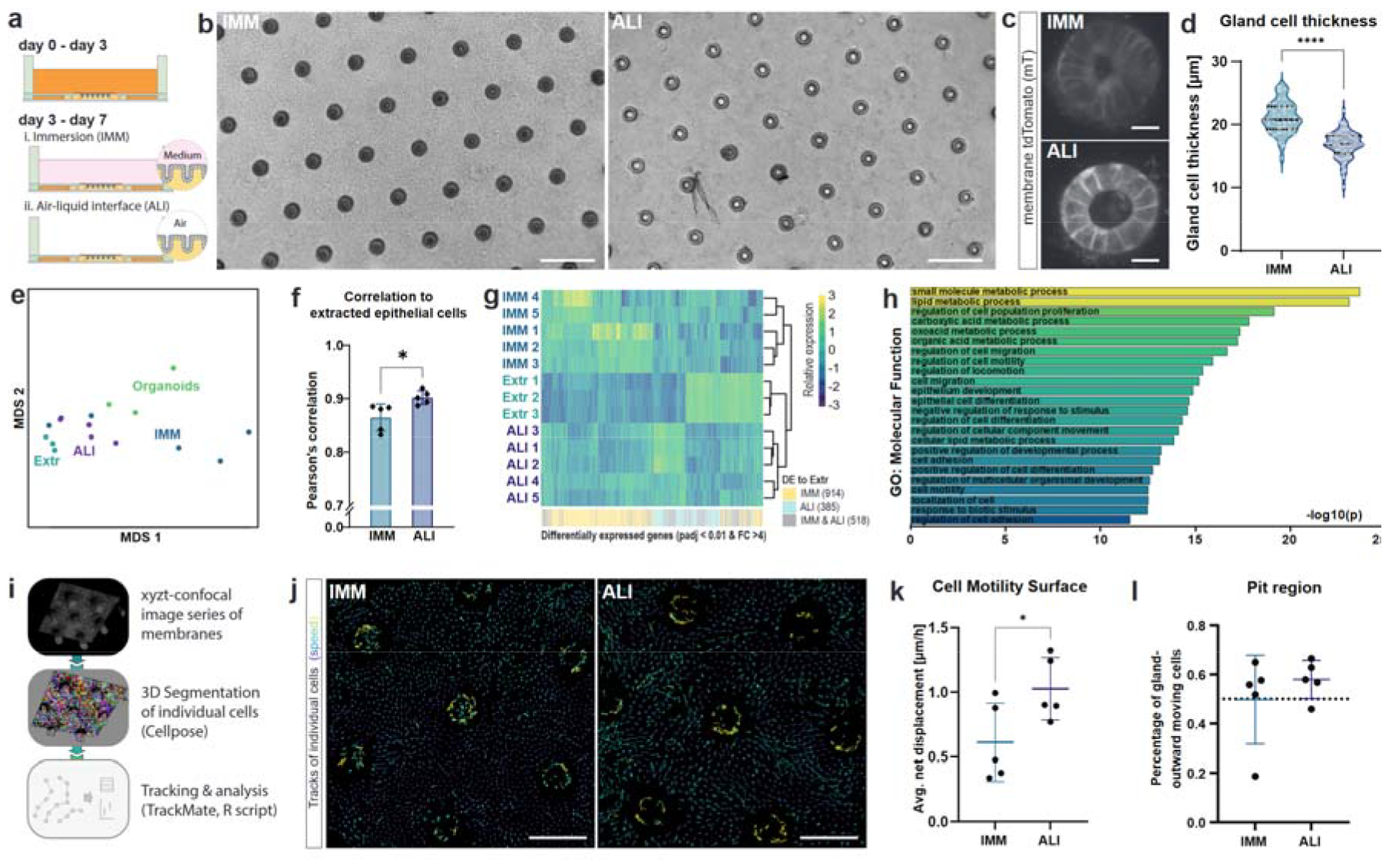
Gastric Transgel organoids in ALI cultures show enhanced transcriptional relevance and increased motility. **a**, Schematics of experimental design. After gastric Transgel organoids were grown for 3 days in expansion medium, the apical side was put to either immersion (IMM) culture (medium with reduced stem cell growth factors) or in air-liquid interface (ALI) culture (apical air). **b-d**, Brightfield (b) and live confocal (c) images of Transgel gastric organoids and quantification of gland cell thickness (d). Each point represents the average cell thickness of one gland. Symbol represents experimental replica (n=6). Horizontal lines show median and quartiles. Scale bars, 200 μm (b), 20 μm (c). **e**, Multidimensional scaling (MDS) plot of RNA sequencing results of samples in IMM and ALI culture and freshly extracted cells (Extr) and Organoids samples as comparison. **f**, Correlation score of RNA sequencing results of Transgel organoids grown in IMM and ALI conditions compared to freshly extracted cells on full transcriptome. Bars indicate mean and error bars SD of 5 independent experiments. **g**, Hierarchical clustering of RNA sequencing results using differentially expressed genes. Indication of genes statistically differentially expressed (DE) in either IMM, ALI or both when compared to freshly extracted epithelial cells. **h**, Enrichment for gene ontology (GO) molecular function terms on genes differently expressed between ALI and IMM Transgel organoids. **i**, Schematics of analysis pipeline. Time-lapse confocal images of ALI and IMM Transgel organoids were acquired and individual cells were identified and tracked over time using Cellpose and Trackmate softwares. **j**, Representative image of tracks of single cells of Transgel organoids in IMM and ALI cultures over 7 hours. Representative image of five independent experiments. Scalebar, 100 μm. **k**, Quantification of movements of cells in the surface regions. Each point represents mean of all cells in one independent experiment, horizonal line represents mean with SD. **l**, Percentage of cells in the proximity of the glands (pit region) that move outwards. Horizontal line represents mean with SD of 5 independent experiments, dotted horizontal line represents expected value of 50% if directionality was random **d,f,k**, Statistical significances from unpaired two-tailed Student *t*-tests are indicated: _*_, p<0.05, _****_, p<0.0001

We further investigated the impact of ALI culture on cell differentiation, by comparing the transcriptome of ALI and IMM Transgel organoids through RNA sequencing. Multidimensional scaling (MDS) and overall correlation analysis showed higher similarity of the transcriptomes of cells in ALI cultures and freshly extracted epithelial cells (**Fig. 3e-f**). Moreover, ALI cultures reduced the number of genes differentially expressed compared to freshly extracted cells from a total of 1’432 genes in IMM cultures to only 903 genes (**Fig. 3g**). Together, this indicates that ALI cultures better recapitulate the gene expression patterns of the real tissue. A gene set enrichment analysis of differentially expressed genes between the two culture methods revealed differences in metabolism, epithelial cell differentiation, and cell motility (**Fig. 3h, Supplementary Fig. S4c**). To validate the observed differences in cell motility, we performed live confocal time-lapse imaging to track individual cells (**Fig. 3i**). Cells in the surface regions of ALI cultures exhibited increased motility compared to those in IMM cultures (**Fig. 3j,k, Supplementary Video 2**). Additionally, cells in proximity to the glands (“pit” regions), but not those further away, displayed a preference for moving away from the glands, resembling the gland-to-surface movement observed in native GI tissues (**Fig. 3l, Supplementary Fig. S4d**)^25^. This observation confirms the preservation of this “conveyor belt” mechanism in our model system.

### Transgel caecal models enable live imaging of syncytial tunnel formation by *T. muris* L1 larvae

Infection studies using classical epithelial organoid cultures are limited by their short lifespan and the lack of apical accessibility imposed by their closed cystic structures. To overcome this problem, microinjections or flipping the polarity of 3D organoids have been used to study viral and bacterial infections^26–28^, but these approaches are not suitable for larger pathogens like multicellular parasites. In addition, they do not allow the modelling of long-term infections nor do they allow the live imaging of the interactions of pathogens with the epithelium^29,30^.

Therefore, we aimed to demonstrate the usefulness of Transgel organoid models for infection studies by infecting caecal models with larvae of the roundworm *T. muris*. During infections *in vivo*, larvae hatching preferably occurs in the caecum, where L1 larvae penetrate the epithelial cell layer and remain solely within those cells, where they create *syncytial tunnels* as they move through the epithelium^31^. Using transwells, we previously developed the first *in vitro* model system for *T. muris* L1 larvae infection and could recapitulate syncytial tunnel formation and study larvae-host cell interactions^14^. However, in addition to the lack of controlled geometrical patterning in that model, the membrane of the transwell obstructed live observations of these host-pathogen interactions.

We added *T. muris* L1 larvae to the apical side of the caecal Transgel organoid model and imaged them using time-lapse microscopy **(Fig. 4a, Supplementary Video 3**). Within 12 hours in ALI cultures, approximately 20% of the larvae became intracellular, while in IMM cultures, the larvae mainly failed to infect. We employed live confocal microscopy using mTmG cells and immunostaining of P43, the most abundant protein of the larvae^32^, on fixed samples to confirm that the larvae indeed penetrated the cells and became intracellular (**Fig. 4b,c, Supplementary Fig. S5a**).

**Fig. 4.**
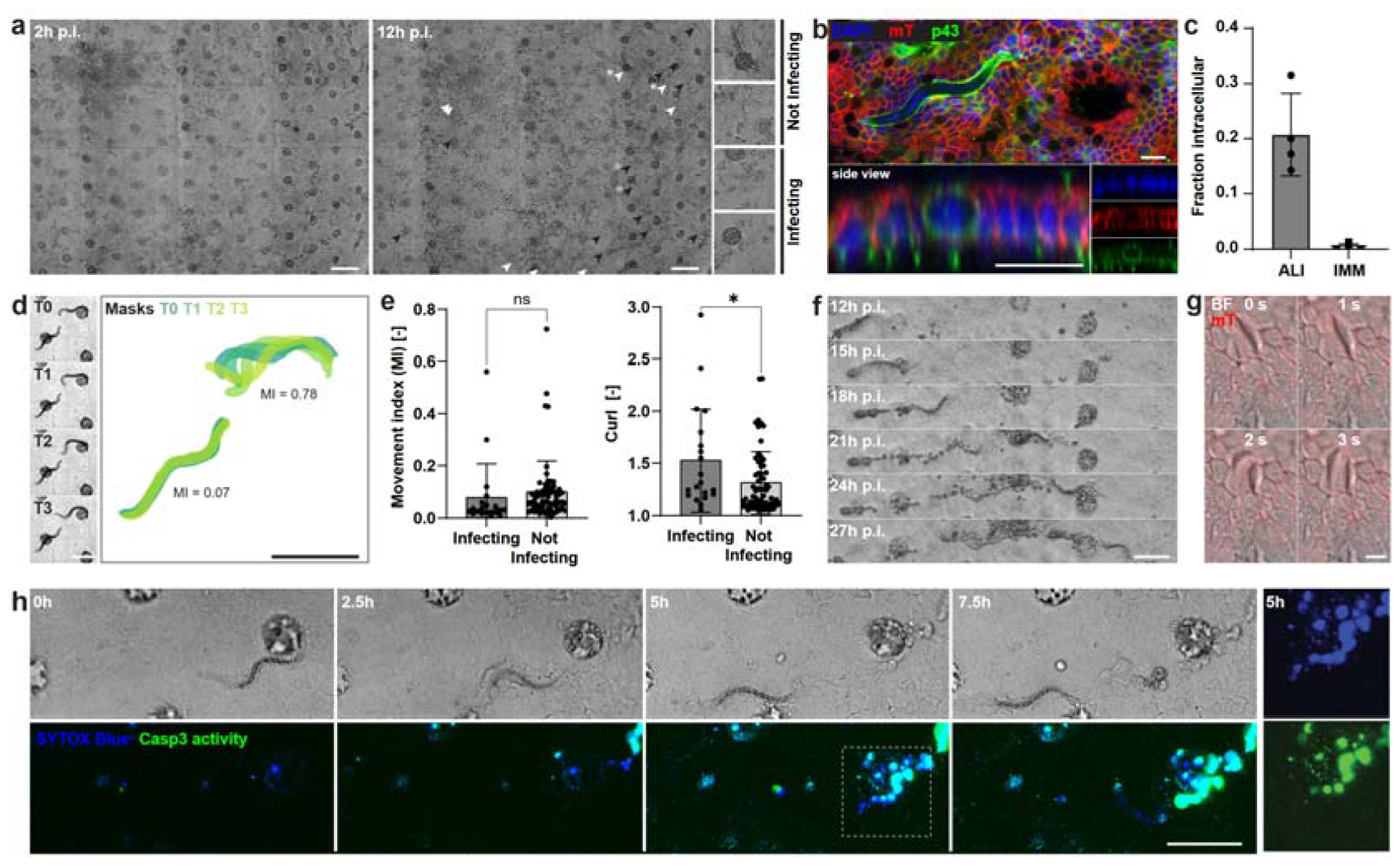
Infection of caecal Transgel organoids with L1 larvae of *T. muris*. **a**, Brightfield overview pictures of caecal Transgel organoids infected for 2 and 12 h with L1 larvae of *T. muris*, in air-liquid-culture (ALI) from 2h p.i. onwards. At 12h p.i. some larvae successfully infected the epithelium and became intracellular (white arrowhead), whereas other didn’t (black arrowheads). Right, insets on example larvae marked with asterisks. Scale bar, 200 μm. **b**, Confocal image of immunofluorescently stained whole-mount sample showing L1 larvae (p43) being intracellular. mT, membrane-tdTomato. Scale bar, 20 μm. **c**, Single L1 larvae were tracked over time using timelapse microscopy. Quantification of proportion of larvae that successfully infected the epithelium and became intracellular in ALI and IMM culture. Each point represents proportion of one independent experiment, bars represent mean with SD. **d**, Movies were acquired at the beginning of infection to assess larval motility and shape. Example image sequence (1 fps) showing two larvae with different motility and the overlay of mask of automated segmentation. Scale bar, 100 μm. **e**, Quantification of larval motility and curl of larvae at the beginning of infection grouped in those that subsequently infect the epithelium and those that don’t. Each point represents one larvae from total four independent replicas, bars represent mean with SD. Statistical significances from unpaired two-tailed Student *t*-tests are indicated: ns, not significative, _*_, p<0.05, **f**, Brightfield image sequence of larvae travelling through epithelial monolayer forming a *syncytial tunnel* and leaving traces of dead cells behind. Scale bar, 100 μm. **g**, High magnification confocal image sequence of intracellular larvae exemplifying the larval head movements. Scale bar, 10 μm. **h**, Confocal image sequence starting 30h p.i. of larvae moving through epithelium forming a tunnel with staining for cell membrane permeability with a nuclear dye (SYTOX Blue) and Caspase 3 (Casp3) activity. Right, single channel inserts of 5h timepoint. Scale bar, 100 μm.

As drug development assays generally measure larval motility, we attempted to assess whether it is a determining factor for successful infection of the epithelium (**Supplementary Fig. S5b**). Immediately after seeding L1 larvae on the epithelial monolayer, we observed considerable heterogeneity in their movements (**Fig. 4d**). Tracking the larvae over time and assessing whether they subsequently became intracellular, we did not detect a significant difference in initial motility between larvae that infected the epithelium and those that did not, indicating that even larvae with low motility can infect the epithelium. However, we found that larvae that infected the epithelium showed a higher level of curl at the beginning of the infection (**Fig. 4e**).

When leaving the infection for longer times, we observed larvae traveling through the epithelial monolayer and forming characteristic *syncytial tunnels* lined with traces of dead cells, similar to what has been previously described (**Fig. 4f**)^14,33^. It appears that larvae typically use their head movements to force through the epithelium (**Fig. 4g, Supplementary Video 4**). Exploiting our highly observable system, we followed single intracellular larvae using confocal microscopy and stained for apoptosis using a dye for caspase 3 (Casp3) activity as well as for membrane disintegration using the membrane-impermeable nuclear dye SYTOX Blue. From cells that were in contact with the larvae, we observed positive signals for dead cells only approximately three to five hours after the larvae passed through the epithelium, indicating the integrity of the membranes during this period. Subsequently, the cells underwent apoptosis, as indicated by the emergence of Casp3 activity (**Fig. 4h, Supplementary Video 4**). *In vivo*, larvae show a preference for moving towards the crypt bottom^33^. We occasionally observed larvae travel to the crypt regions of the model system, where they seemingly became stuck (**Supplementary Fig. S5c**).

To our knowledge, this is the first *in vitro* model that allows live observation of *T. muris* L1 larvae traveling through a caecal epithelial monolayer. We found that larvae infecting the cells create tunnels to move through the epithelium, similar to what occurs *in vivo*, and we characterized the cell death that typically occurs by activation of an apoptotic pathway subsequent to membrane disintegration.

## Discussion

In this study, we developed Transgels, a platform for generating bilaterally accessible 3D tissue models of GI epithelia, which demonstrated high physiological relevance and allowed for detailed observations of cellular behaviors. As a consequence of the gained accessibility, we could mimic basal stem cell niche GF delivery, which led to proliferative and differentiation patterns consistent with native tissues. We were also able to remove the apically accumulating dead cells that are constantly being shed, allowing for homeostatic conditions.

Transgels use a collagen-matrigel hydrogel as the growth substrate, which provides *in vivo-*like cell attachment sites and physiological stiffness that support stem-cell maintenance and function^19,34^. Moreover, the hydrogel surface’s geometry can be pre-defined to shape the epithelium to match the tissue of origin. This ultimately makes the system physiologically relevant, as shown by a high degree of transcriptional correlation with 3D organoids and epithelial cells freshly extracted from the tissues.

The use of ALI cultures further enhances the transcriptional accuracy of the gastric epithelial model. Compared to conventional 3D models, one of the key advantages of the Transgel organoids is their excellent observability, which let us visualize and track individual cells in real-time. This allowed us to suggest that this increased physiological relevance of ALI cultures is associated with increased cellular motility.

We also observed gland-to-surface movement similar to what has been indirectly shown in native tissues. This suggests that the Transgel organoid system recapitulates the dynamic nature of the gastric epithelium, which is characterized by continuous cell turnover and migration from the stem cell compartments to the surface.

We finally demonstrated the utility of the Transgel organoid system for infection studies by infecting the caecal model with *T. muris* L1 larvae. We demonstrated the capability of the model to recapitulate larval epithelial cell invasion and tunnel formation, key events of early *T. muris* infections. By tracking individual larvae and analyzing the cellular responses, such as membrane disintegration and apoptotic pathways, we gained insights into the interactions between the larvae and the host epithelium. Interestingly, even cells directly penetrated by the larvae did not die immediately but remained intact, preserving epithelial function for several hours. Additionally, we found that larval motility was not a determining factor for successful infection, as even larvae with low motility succeeded in infecting the epithelium. This suggests that other unknown factors play a crucial role in the infection process. Unlike previous caecal epithelial models that were based on membranes^14^, our study provided an opportunity for larvae to penetrate the epithelial barrier and invade the hydrogel. However, we did not observe such invasion, confirming that the formation of tunnels within the epithelial cells is an inherent property of the interaction between larvae and epithelial cells, rather than a result of larvae actively avoiding underlaying tissue components present *in vivo*, such as for instance cells of the immune system. We thus recapitulate critical host epithelial cell – parasite interactions that determine the tropism of the larvae.

In conclusion, the Transgel-based 3D tissue models provide a valuable platform for studying the physiology and pathology of GI epithelia. In contrast to traditional 3D organoids, the models are conveniently accessible on both the basal and apical sides independently and allow for real-time observation of cellular behaviors. These features enable the modeling and direct observation of long-term infections with parasites such as *T. muris*. We believe that observable and accessible 3D organoid systems, such as the Transgel platform presented in this study, hold great promise for advancing our understanding of epithelial biology and infectious diseases.

## Data Availability

The main data supporting the results in this study are available within the paper and its Supplementary Information. Source RNA-seq data has been deposited to the Gene Expression Omnibus (GEO) public repository with the accession code GSE241012.

## Code Availability

Custom analysis code is available on request.

## Author contributions

M.H. and M.P.L. conceived the study, designed experiments, interpreted data and wrote the manuscript. M.H. conducted all experiments and analysis. M.A.D-C. proposed and supervised *T. muris* infection experiments, provided *T. muris* eggs, contributed to data interpretation and edited the manuscript.

## Acknowledgements

We thank Dr. Kunal Sharma from the Laboratory of Prof. John McKinney (EPFL) and Dr. Alexandre Widmer from the EPFL Organ/Tissue Sharing Program for providing cadavers of mice and Prof Richard Grencis (University of Manchester) for the P43 antibody. We thank Dr. Mikhail Nikolaev for help producing the microfabricated molds for stamps, Dr. Sophia Li for help generating colonic organoids, Antonius Chrisnandy, Bilge Sen Elci and Tania Hübscher for valuable discussions and Julia Prébandier, Lucie Tillard and Charlotte Tolley for administrative and technical support. We acknowledge support from CMi, BIOP, GECF, FCCF and HCF EPFL core facilities. This work was funded by the National Center of Competence in Research Bio-Inspired Materials, the EU Horizon 2020 research program INTENS (www.intens.info; no. 668294-2) and the Swiss National Science Foundation research grant no. 310030_179447.

## Competing interests

The other authors declare no competing interests.

## Additional information

### Methods

#### Microfabrication of Transgel devices

Molds for Transgel devices were fabricated using conventional photolithography. Briefly, two 200 µm layers (total 400 µm) of SU-8 photoresist (Gersteltec, GM 1075) were spin-coated on a 150 mm silicon wafer, soft baked and UV-exposed applying a chrome mask. After baking and development steps, the mold was coated with chlorotrimethylsilane (Sigma-Aldrich, 33014) by vapor deposition. 10.5g of polydimethylsiloxane (PDMS, Sylgard 184) with a curing agent ratio of 1:10 was poured on the wafer, yielding a total height of 800 µm. A second layer of PDMS (h = 1 cm) as superstructure containing a central well (d = 12 mm) was O_2_-plasma-bonded on top of the microfabricated layer. Holes for the basal side reservoirs (d = 4 mm), gel compartment opening (d = 4 mm) and hydrogel loading port (d = 1.5 mm) were punched using biopsy punchers (Miltex) and the total structure was O_2_-plasma-bonded to a glass coverslip.

For hydrogel loading, either flat or microfabricated PDMS stamps with the desired geometries were applied on the center of Transgel devices. Geometries as previously reported were used with adaptions for stomach (gland depth = 300 µm) and caecum (crypt depth = 100 µm) systems^1^.

5 mg/ml Collagen type I (Koken, KKN-IAC-50) was neutralized (1:4 addition of 5X DMEM supplemented with 50 mM NaHCO_3_ (Thermo Scientific, J63025AK)) and mixed with Matrigel (Corning, growth factor reduced, phenol red-free formulation) at a ratio 4:1 on ice and loaded to the Transgel devices using the hydrogel loading port (Fig. 1a) and let to gelate at 37°C for 30 min. For the intestinal model with a geometry with villus domains, 6 mg/ml Collagen Type I (Advanced Biomatrix, 5225) was used instead.

After stamp removal, hydrogels were washed and kept for maximal two weeks in 1X phosphate-buffered saline pH 7.4 (PBS, Gibco, 16210064) at 4°C until use. Before seeding cells, hydrogels were equilibrized for > 4h with growth medium.

#### Diffusion modelling and experimental validation

Diffusion from basal side reservoir through gels was modelled in COMSOL Multiphysics. Diffusion coefficient in collagen hydrogels was estimated from ref.^2^. A model growth factor protein with 40 kDa molecular weight and 3 nm hydrodynamic radius was used. Experimentally, 1 mg/ml fluorescein isothiocyanate (FITC)-labelled 40 kDa dextran was added to the medium on the basal side and diffusion through the gel was observed using confocal microscopy over time. To validate epithelial barrier integrity and the ability to control both sides of the epithelium independently, either Rhodamine B isothiocyanate (RITC)- or FITC-labelled dextran was added on the apical and basal side media of a grown flat epithelial monolayer, respectively, and confocal z-stack images were taken 24h after medium replenishment. Orthogonal (“xz”) average intensity projection are displayed.

#### Mice

Gastrointestinal organs of wild-type or Gt(ROSA)26Sor tm4(ACTB-tdTomato,-EGFP)Luo (“mT/mG”) C57BL/6 mice cadavers were obtained through EPFL’s internal organ/tissue sharing program or from McKinney Laboratory, EPFL, following animal experimentation protocols prescribed by EPFL, in compliance with local animal welfare laws, guidelines and policies.

#### L-WRN-Conditioned medium

Triple Wnt/R-Spondin/Noggin (WRN)-conditioned medium was produced using L-WRN cells (ATCC, CRL-3276) according a modified version of a previously published protocol^3^. In brief, L-WRN cells were grown until confluency Advanced DMEM/F-12 (Gibco, 12634010) supplemented with 10 mM HEPES (Gibco, 15630080), 1X GlutaMAX (Gibco, 35050061), 20% FBS (Gibco, 10270106) and 50 U/ml Penicillin-Streptomycin (Gibco, 15070063). After reaching confluency, medium was replaced and collected every 24 hours for 9 days. Collections of three consecutive days were pooled and diluted 1:1 in medium without FBS (final FBS concentration: 10%), sterile filtered, and stored at -20°C.

#### Organoid generation and culture

Gastrointestinal epithelial cells were extracted from the antrum, small intestine, caecum and colon regions following previously published protocols^4–7^. Obtained cell aggregates were embedded in 25 µl Matrigel drops in 24-well plates and cultured in tissue-specific expansion media (Supplementary Table 1). For passaging, organoids were collected from Matrigel in ice-cold adv. DMEM/F-12 and mechanically disrupted using a glass Pasteur pipette. Fragments were diluted in fresh Matrigel at a split ratio between 1:2 and 1:6, depending on density. To compare organoids to transgels and freshly extracted epithelial cells, organoids were grown for three days in expansion medium, and then for two additional days (Stomach: four days) in differentiation medium. Organoids between passage numbers 5 and 15 were used for experiments.

#### Cell seeding and culture on Transgel devices

Organoids were collected in ice-cold adv. DMEM/F-12 and dissociated using TrypLE Express (Gibco, 12605028) for 12 min at 37°C, with mechanical disruption using a P1000 pipette every 4 minutes. Cells were diluted in adv DMEM/F-12 and strained using a 70 µm strainer and spun down. Medium was removed from the Transgel devices and 8 µl of a 12.5 mio cells/ml single cell solution was added on top of the hydrogel, avoiding the cells to spread on the PDMS. Cells were let to sediment (30 min, 37°C) before adding expansion medium. On day 3, the apical side expansion medium was replaced with differentiation medium containing reduced concentration of growth factors (Supplementary Table 1), and kept for an additional two days (Stomach: four days) in culture. Where indicated, the medium was completely removed (air liquid interface [ALI] cultures). Medium was replaced every two days.

#### Cell density, viability and organoid forming assays

From fluorescent images of the cell membranes, cell density was assessed by segmenting cells with CellProfiler (v4.2.5, ref^8^). Resulting densities of three regions per sample were averaged.

From Transgel devices, single cells were obtained by digesting the hydrogel with the cells in 10’000 U/ml Collagenase Type 1 (Gibco, 17100017) for 12 min at 37°C and by subsequently dissociating the cells with TrypLE Express for 10 min at 37°C. From organoids, single cells were obtained as described above. A subset of cells was stained with 10 µg/ml Hoechst (Thermo Scientific, 62249) and 1 µM DRAQ7 (BioLegend, 424001) for 20 min at 37°C and imaged subsequently. Total cells were counted using Hoechst channel image and for each cell the DRAQ7 intensity was measured. Cells were classified as being dead when their DRAQ7 signal exceeded the background signal by at least 3x it’s standard deviation. Similarly obtained single cell solutions were replated at 0.5 mio cells/ml in Matrigel and cultured in expansion medium. Organoid forming efficiency was assessed as the number of organoids that grew after 3d divided by the initial number of cells per region. Values from three regions (10X stack images) per independent replica were averaged.

#### Immunofluorescence and histological staining

Collagen gels with cells were removed from the Transgel devices and fixed in 4% paraformaldehyde (ABCR, AB351601) for 30 min at room temperature (RT). Samples were then washed with PBS three times for 1h at RT and incubated in 50% Cryomatrix embedding resin (Epredia, 6769006) in PBS at 4°C overnight. Samples were then embedded in 100% Cryomatrix in a plastic specimen holder and frozen on dry ice. 10 µm-thin cryosections were obtained using a Leica CM3050 S cryostat cutting at -20°C and immobilized on microscopy slides. Sections were permeabilized with 0.2% Triton X-100 (Sigma-Aldrich, X100) in PBS (15 min, RT) and blocked in 10% goat serum in PBS containing 0.01% Triton X-100 (30 min, RT). Samples were incubated with primary antibodies (Supplementary Table 2) diluted in blocking buffer (overnight, 4°C) and washed with blocking buffer three times for 30 min. Samples were then incubated with secondary antibodies, DAPI, and fluorophore-conjugated phalloidins (Supplementary Table 2) for 2h at RT and washed three times with PBS. Whole mount stainings were performed similarly with longer incubation times: Fixing, 1h; Permeabilization, 1h; Blocking, 3h; Primary and secondary antibodies, overnight; Washes, 3x3h plus overnight.

For staining of mucins, samples were fixed in methacarn fixative (60%v/v methanol, 30% chloroform, 10% acetic acid) for 1h at RT. Samples were washed in PBS and cryosections were obtained as above. Sections were stained with Alcian blue (AB) at pH 2.5.

#### Microscopy and image processing

Brightfield and red fluorescent imaging of living samples was performed using a Nikon Eclipse Ti2 inverted microscope system equipped a 555 nm filter cube, and a DS-Qi2 camera using 4x/0.20NA and 10x/0.30NA objectives. Confocal imaging of living samples was performed using a Leica SP8 inverted microscope system equipped with a supercontinuum laser (range 470 nm-670 nm) and hybrid photon counting detectors (HyD), using a 2.4 mm WD HC FLUOTAR 25x/0.95NA water-immersion objective. Immunofluorescently stained sections were imaged with an upright Leica SP8 microscope system equipped with 405 nm, 488 nm, 552 nm and 638 nm solid state lasers and HyD detectors using HC PL APO 20x/0.75NA and HC PL APO 63x/1.40NA objectives. AB stained sections were imaged on an upright Leica DM5500 system equipped with LED illumination and a DMC 2900 Color camera, using a HC PL FLUOTAR 20x/0.7NA objective and white balance correction.

Image processing was performed using FIJI ImageJ^9^ (v1.54d) using standard contrast and intensity level adjustments, for noise filtering, for drift and tilt corrections, as well as for generating z-maximum intensity and orthogonal projections. Extended depth of field (EDF) images were generated with LAS X software (Leica).

#### RNA extraction, library preparation and sequencing

Cells were collected via collagenase digestion (Transgel hydrogels) or in ice-cold adv DMEM/F12 (organoids) as described above. Control samples from fresh tissues were obtained following previously published protocols^4,6,7^. Purity of fresh epithelial cell extractions was assessed by flow cytometry. Therefore, obtained single cells were stained for EpCAM and with DAPI (Supplementary Table 2) in PBS containing 10% FBS, 2 mM EDTA (Invitrogen, 15575020) for 10 minutes on ice and washed twice in the same buffer. Flow cytometry was performed on an LSRFortessa instrument (BD Bioscience) gating on live, single cells. RNA was extracted using RNeasy Micro kit (Qiagen, 74004) and quantified by Qubit Fluorometer (Invitrogen). Libraries for multiplexed illumina sequencing were generated following the QuantSeq 3’ mRNA-Seq Library Prep Kit FWD (Lexogen) with 100 ng RNA input and 17 PCR cycles. Quantity and quality of the libraries were assessed with Qubit Fluorometer and DNF-474 HS NGS Fragment Analyzer (Aligent). Sequencing was performed on a NextSeq500 (Illumina) machine with >8.7 mio requested reads per sample and raw count matrices were generated by STAR v2.7.0e. Analysis was performed in the RStudio environment following edgeR analysis pipeline^10^. Briefly, raw counts of n = 3 independent replicas were normalized, logarithmized and lowly expressed genes were removed. Tissue signature genes were defined as genes differently expressed (DEG) using a genewise negative binomial generalized linear model with quasi-likelihood test (FDR<0.001) and used for correlation (Pearson’s) analysis. For analysis of gastric Transgel organoids, sample size was increased to n = 5. Gene ontology analysis was performed using *goana* (limma) function on DEG (p-value<0.01, log fold change < 4), between ALI and IMM culture samples and ontology terms were filtered by the number of total genes associated (700<N<1800) to unselect too specific and too general terms.

#### Cell motility analysis

Confocal stack images of fluorescent membranes were acquired every hour. For cell segmentation, a customized model was trained in cellpose 2.0^11^ using multiple single xy and xz planes from the data starting from the LC1 model with loops of manual corrections. The model was applied to the 3D images of each timepoint. Output label images were analyzed using FIJI TrackMate plugin using Kalman tracker^12^. 3D output paths were analyzed in R software to analyze directionality and average speed and to group by experiment. Path were classified by the distance *d* to the glands and relative *z* position (surface: *z* = 0) in *gland* (*z* < -30 µm), *pit* (*d* < 40 µm), and *surface* (*d* > 40 µm) regions.

#### Infection with *T. muris* first-stage (L1) larvae

Embryonated *T. muris* eggs were hatched as previously described^13^. 50-100 purified L1 larvae were added to 5-day-old transgel caecum organoids and let to settle. 2h p.i. medium was completely removed to induce air-liquid interface conditions unless indicated otherwise. Typically, 10-70 larvae remained on the epithelium during this process. Samples were live-stained with 5 µM NucView 488 Caspase 3 dye (Sigma-Aldrich, SCT101) and 3 µM SYTOX Blue membrane-impermeable nuclear dye (Invitrogen, D15106) 20 minutes before imaging.

To assess larval motility at the beginning of infection, larvae imaged for 5 s at 3 fps were segmented with Ilastik software^14^ (v1.4.0,) using a two-step pixel and object classification algorithm. The movement index was calculated as the ratio between the area of the XOR images between two consecutive masks and the average size these two masks (Supplementary Fig. 5b) using FIJI. The median value per larvae was reported. The average curl of each larvae was assessed by the ratio between the length of the skeleton of the mask (larvae length) and the eucledian distance between the extremities. The median per larvae over 5 images at 1 fps was reported.

#### Reporting Summary

Further information on research design and methods is available in the Nature Research

**Supplementary Table 1:**
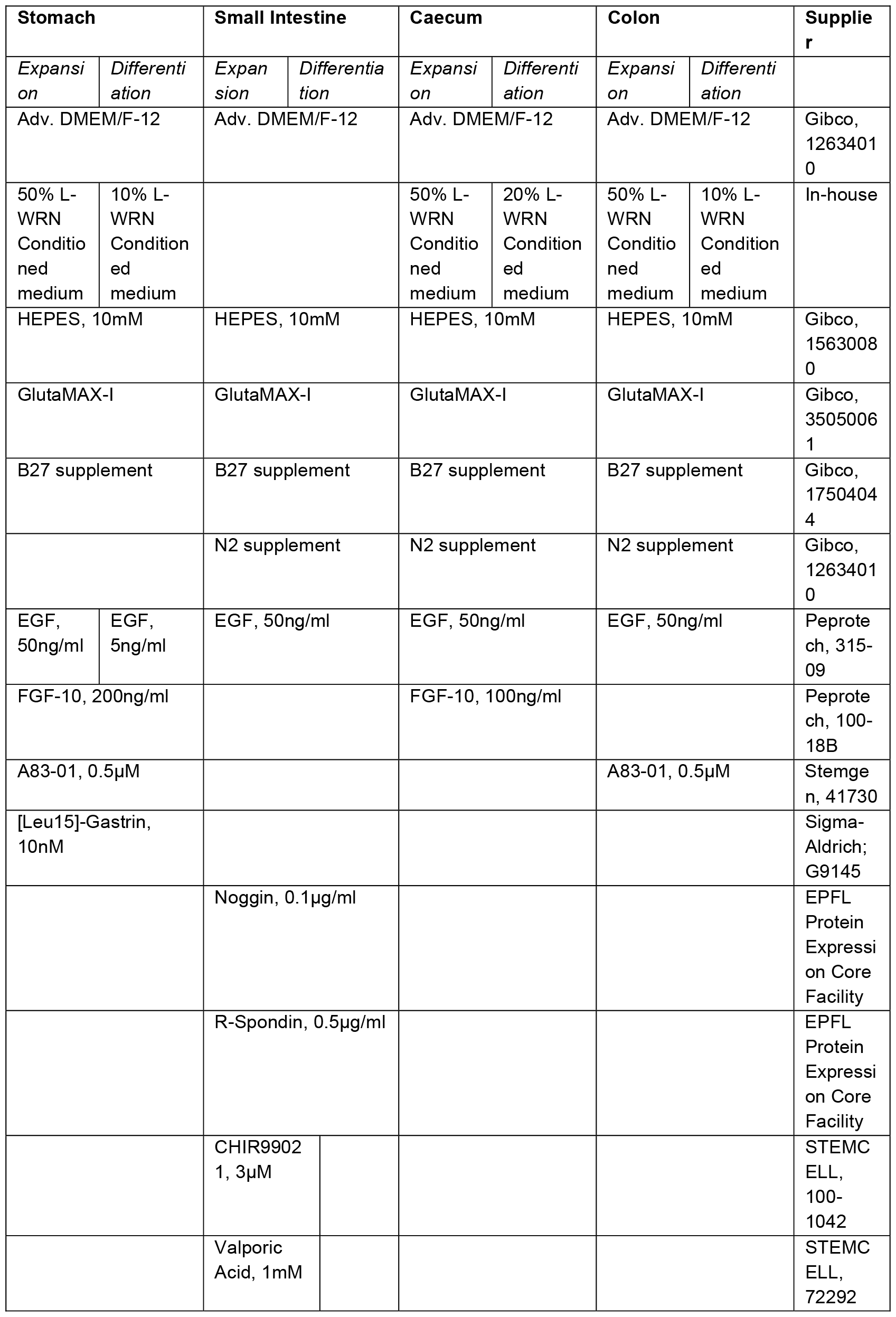
Media formulations for organoid and Transgel organoid culture.

**Supplementary Table 2:** Antibodies used for Immunofluorescent stainings (IF) and flow cytometry (FC). DAPI, 4′,6-diamidino-2-phenylindole.

| Target | Conjugate | Dilution | Supplier |
| --- | --- | --- | --- |
| **Primary Antibodies** |  |  |  |
| Sox9 |  | 1:200 | Abcam, ab185966 |
| Ki67 |  | 1:100 | BD Biosciences, 550609 |
| Itga6 |  | 1:100 | Sigma Aldrich, MAB1378 |
| P43 |  | 1:100 | Prof Richard Grencis, University of Manchester |
| CD326 (EpCAM) | APC | 1:800 (FC) | Invitrogen, 17-5791-82 |
| **Secondary Antibodies** |  |  |  |
| Mouse IgG | Alexa Fluor 488 | 1:400 | Invitrogen, A-11029 |
| Rabbit IgG | Alexa Fluor 647 | 1:400 | Invitrogen, A-31573 |
| Rat IgG | Alexa Fluor 647 | 1:400 | Invitrogen, A-21247 |
| **Others** |  |  |  |
| Phalloidin | Alexa Fluor 546 | 1:100 | Invitrogen, A22283 |
| DAPI |  | 2.5µg/ml (IF),<br>0.5µg/ml (FC) | Sigma-Aldrich, D9542 |

## Supplementary Figures

**Supplementary Fig. S1.**
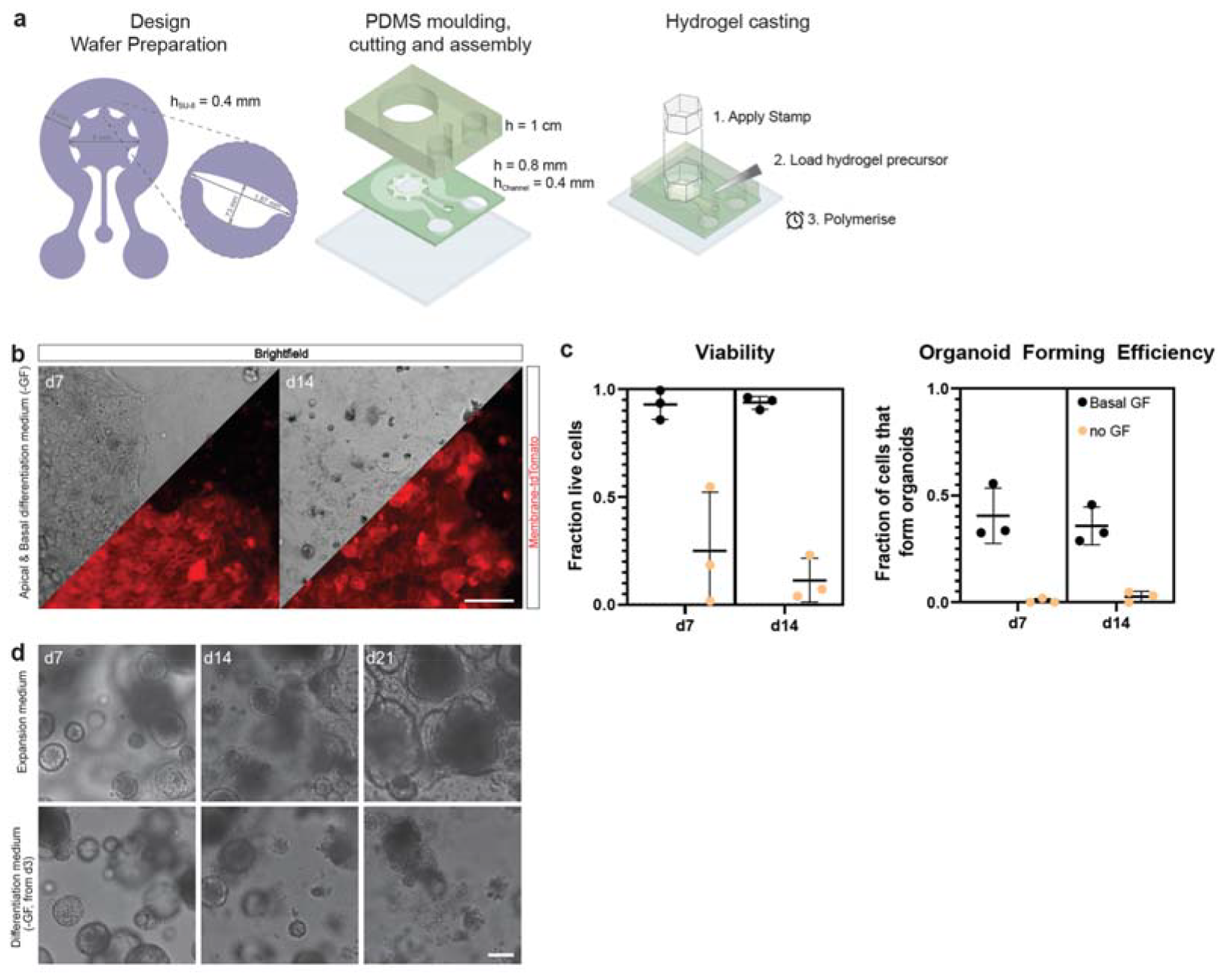
Fabrication of Transgel device. **a**, Design and fabrication of Transgel devices. First, a microfabricated wafer with SU-8 topology was prepared, used for the first layer of Transgel device. This layer is oxygen-plasma-bonded to glass, and a well-like superstructure is bonded onto it. To cast the hydrogel, a polydimethylsiloxane (PDMS) stamp is temporarily applied at the opening, and hydrogel precursor solution is loaded via the hydrogel access port. **b,c**, Control to Fig. 1g-i, Brightfield picture and assessment of cell viability and organoid forming efficiency of day 7 and day 14 of gastric epithelial cells when removing of stem cell growth factors (GFs) also from the basal side medium from day 3 on. Epithelium maintenance is impaired, showing that basally delivered GFs are necessary for epithelium maintenance. c, Cells were collected from Transgel hydrogels and analyzed for viability and capacity to form organoids. Mean and SD from three independent experiments are shown. Scale bar, 100 μm. **d**, Brightfield image sequence of gastric organoids in expansion (with GFs) and differentiation (without GF), related to Fig. 1i. Scale bar, 100 μm.

**Supplementary Fig. S2.**
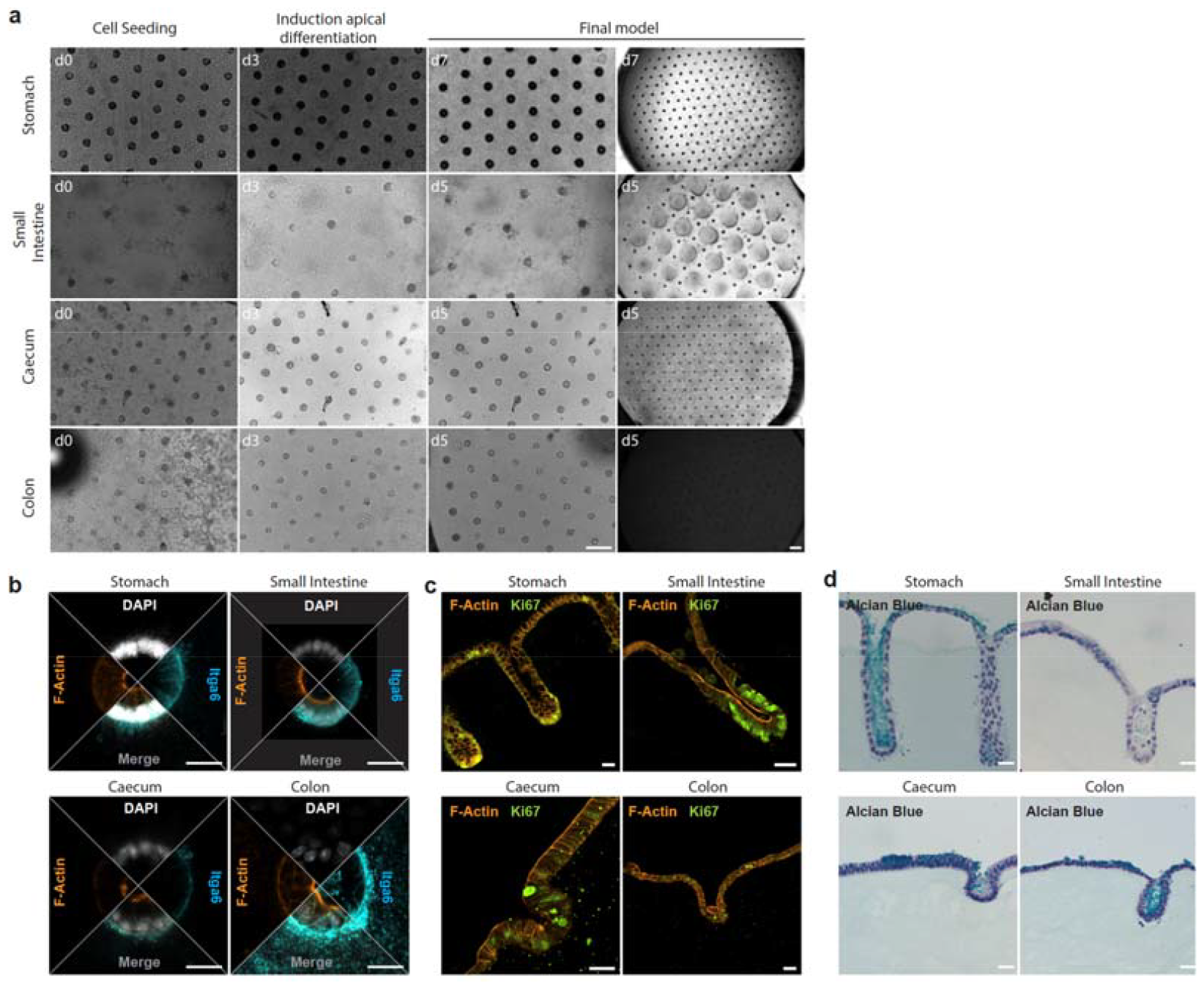
Development and characterization of Transgel organoids. **a**, Brightfield image sequence of Transgel organoids growth. Cells from tissue-specific organoids were seeded on hydrogel scaffolds and grown for 3 days in expansion medium (with stem cell growth factors (GF)) to form an epithelial monolayer. From then on, the concentration of GF on the apical side was reduced (see methods) and cells were cultured for an additional 4 (stomach) or 2 (others) days, until the final model was achieved. Scale bars, 200 μm. **b**, Confocal image of Transgel organoids at crypt/glands, immunofluorescently stained for Itga6 and F-actin. Scale bars, 20 μm. **c**, Immunofluorescent staining for the proliferation marker Ki-67 and F-actin of cryosections of Transgel organoids. Scalebars, 20 μm. **d**, Alcian blue staining on cryosections of Transgel organoids. Scale bars, 20 μm.

**Supplementary Fig. S3.**
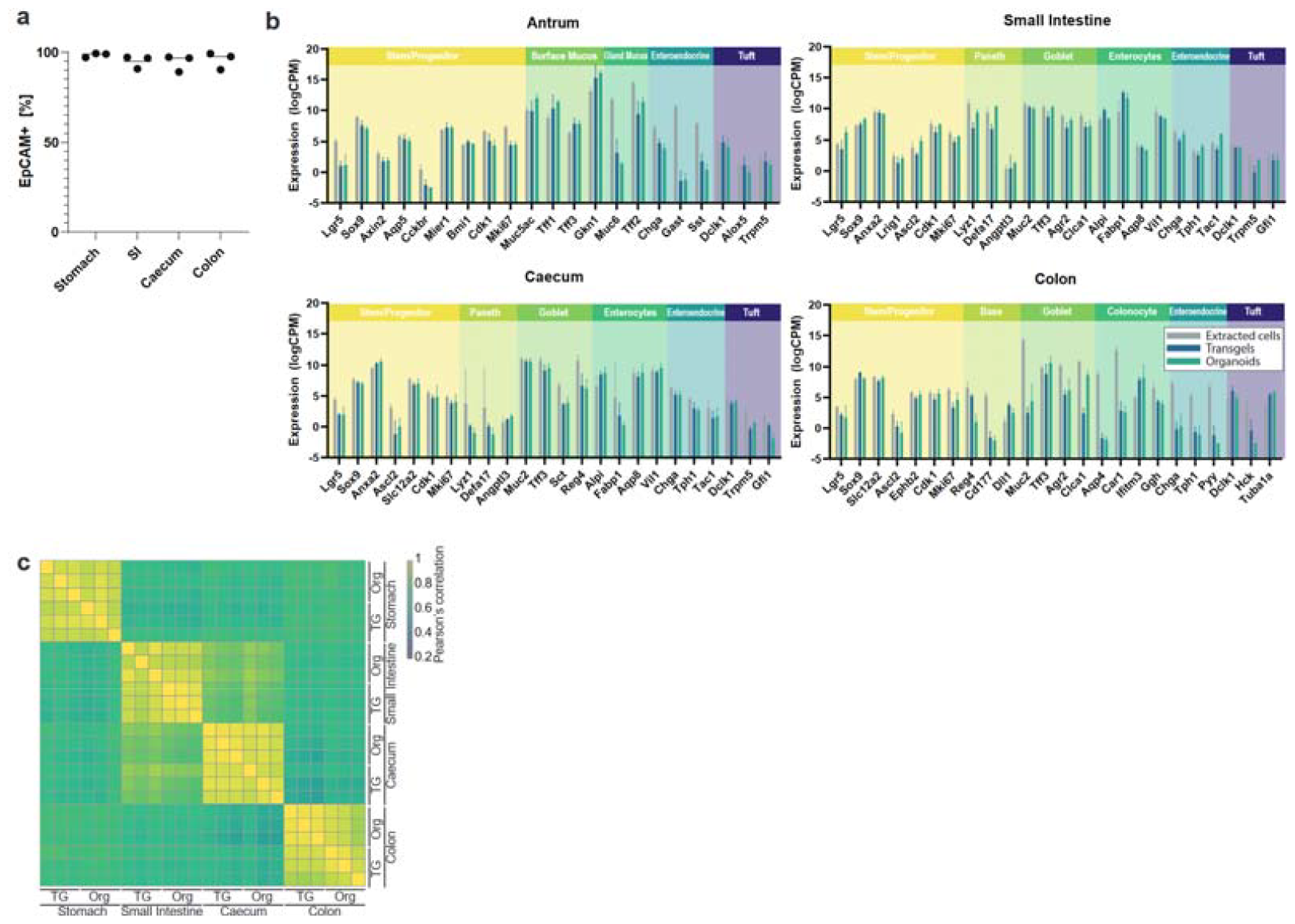
Transcriptional comparison between organoids grown in 3D and Transgel cultures. **a**, Purity assessment of freshly extracted epithelial cells used as reference for RNA sequencing experiment by flow cytometry. Fraction of EpCAM+ cells gated for cells, single cells, live cells. Each point represents one independent extraction from a different mouse, horizontal bar represents mean value. **b**, Expression levels (mean and SD of three independent experiments) of genes of interest in Transgel organoids, 3D organoids and freshly extracted epithelial cells. **c**, Correlation score of all *in vitro* models (Transgel organoids [TG] and 3D organoids [Org]) show high transcriptional correlation between the two culture methods.

**Supplementary Fig. S4.**
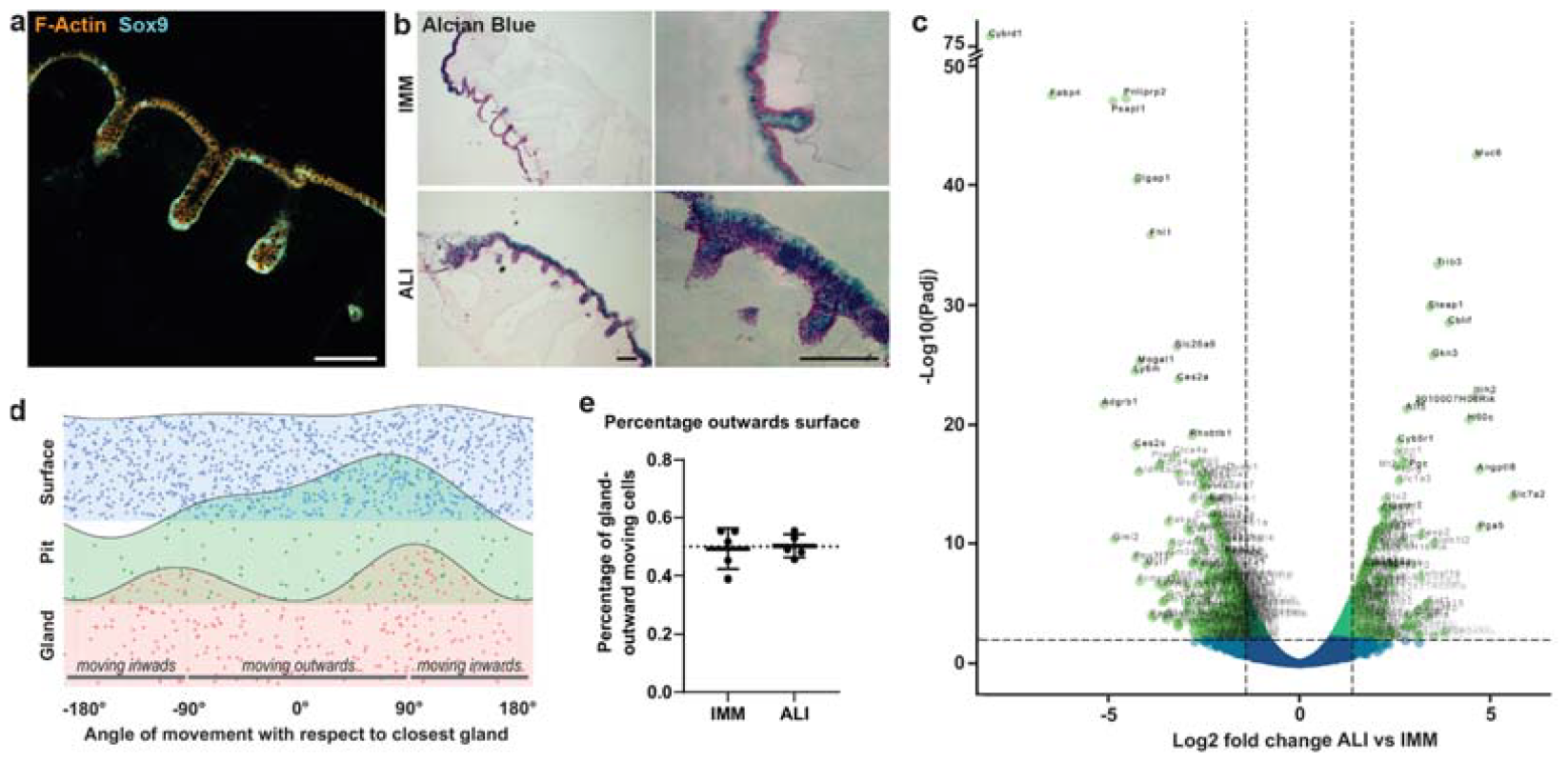
Characterization and comparison of Transgel organoids cultured in air-liquid interface (ALI) and immersion (IMM) culture. **a**, Immunofluorescent staining of Sox9 and F-actin of gastric Transgel organoid in ALI culture. Scale bar, 100 μm. **b**, Alcian Blue staining of gastric Transgel organoids in IMM and ALI culture. Scale bars, 100 μm. **c**, Volcano plot of transcriptional comparison between ALI and IMM culture. Related to Fig. 3h. **d**, Representative distribution of angle of movement with respect to closest gland per cell, categorized by location of cell. **e**, Percentage of cells in the surface regions that move outwards from glands. Each point represents one independent experiment and horizontal line represents mean with SD. Dotted horizontal line represents expected value of 50%.

**Supplementary Fig. S5.**
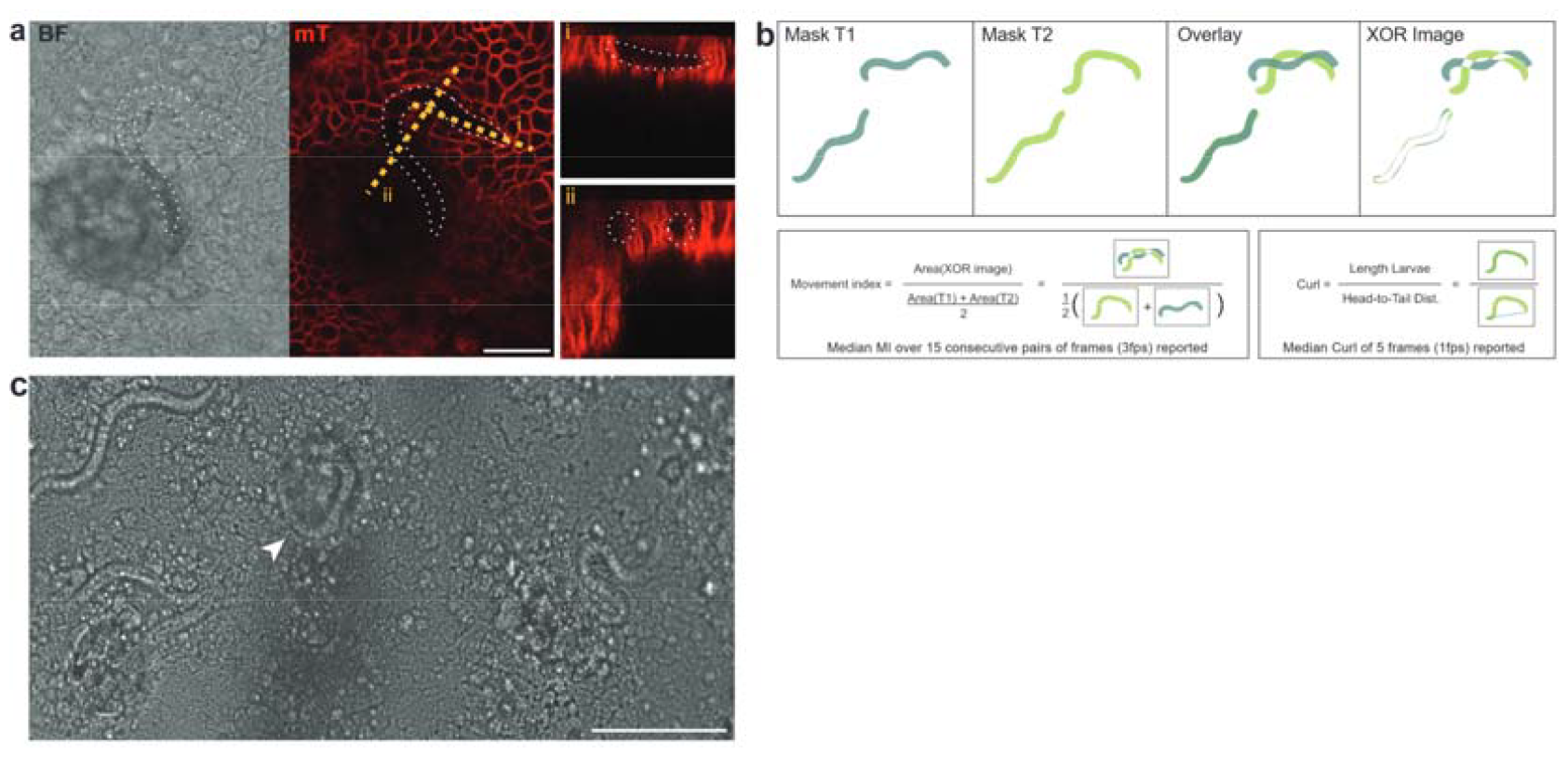
Extended data to *T. muris* infection on caecal Transgel organoids. **a**, Confocal image and side projections (right) of live caecal Transgel organoids with intracellular L1 *T. muris* larvae. Scale bar, 20 μm. **b**, Analysis strategy of movement index and curl of larvae. Related to Fig. 4d,e. **c**, Example of larvae localized in the crypt region. Scale bar, 100 μm.

## Supplementary Videos

**Supplementary Video 1:** Animated confocal live-acquired stack images of stomach, small intestine, caecum and colon Transgel organoids. Brightfield and membrane-tdTomato. Related to Fig. 2c.

**Supplementary Video 2:** Cellular motility of gastric Transgel organoids in immersion and air-liquid interface cultures. Z-coded confocal time lapse images of membrane-tdTomato cells and single slice images at surface and glands as well as 3D reconstruction. Related to Fig. 3j.

**Supplementary Video 3:** Infection of T. murs larvae on caecal Transgel organoids. Overview timelapse images of first 18h of infection. Magnification movie 12h p.i. (live) of larvae at different stages of infection. Some successfully invaded the epithelial monolayer and became intracellular. Animated live-acquired stack image of intracellular larvae, brightfield and membrane-tdTomato channels. Related to Fig. 4a-e.

**Supplementary Video 4:** Trichuris muris larvae moving through epithelium by tunnel formation. Live observation of single larvae by brightfield timelapse imaging. Confocal imaging with staining for dead cells (SYTOX Blue) and apoptosis (Casp3). High magnification movie of larval head movements. Related to Fig. 4f-h.

## Notes

### Competing Interest Statement

The authors have declared no competing interest.

